# ProT-VAE: Protein Transformer Variational AutoEncoder for Functional Protein Design

**DOI:** 10.1101/2023.01.23.525232

**Authors:** Emre Sevgen, Joshua Moller, Adrian Lange, John Parker, Sean Quigley, Jeff Mayer, Poonam Srivastava, Sitaram Gayatri, David Hosfield, Maria Korshunova, Micha Livne, Michelle Gill, Rama Ranganathan, Anthony B. Costa, Andrew L. Ferguson

## Abstract

The data-driven design of protein sequences with desired function is challenged by the absence of good theoretical models for the sequence-function mapping and the vast size of protein sequence space. Deep generative models have demonstrated success in learning the sequence to function relationship over natural training data and sampling from this distribution to design synthetic sequences with engineered functionality. We introduce a deep generative model termed the Protein Transformer Variational AutoEncoder (ProT-VAE) that furnishes an accurate, generative, fast, and transferable model of the sequence-function relationship for data-driven protein engineering by blending the merits of variational autoencoders to learn interpretable, low-dimensional latent embeddings and fully generative decoding for conditional sequence design with the expressive, alignment-free featurization offered by transformers. The model sandwiches a lightweight, task-specific variational autoencoder between generic, pre-trained transformer encoder and decoder stacks to admit alignment-free training in an unsupervised or semi-supervised fashion, and interpretable low-dimensional latent spaces that facilitate understanding, optimization, and generative design of functional synthetic sequences. We implement the model using NVIDIA’s BioNeMo framework and validate its performance in retrospective functional prediction and prospective design of novel protein sequences subjected to experimental synthesis and testing. The ProT-VAE latent space exposes ancestral and functional relationships that enable conditional generation of novel sequences with high functionality and substantial sequence diversity. We anticipate that the model can offer an extensible and generic platform for machine learning-guided directed evolution campaigns for the data-driven design of novel synthetic proteins with “super-natural” function.

## 1 Introduction

Proteins are molecular machines that are the workhorses of biology. Natural proteins have evolved under natural selection to perform diverse biological functionalities, including enzymatic catalysis, molecular transport, cellular signaling, and immunological surveillance. The ability to design synthetic sequences with engineered functionalities is a long-standing goal of synthetic biology with enormous potential in multiple fields including medicine, public health, biochemical engineering, and clean energy. Protein sequence space is almost unimaginably vast: restricting ourselves to just the 20 natural amino acids, there are ~10^260^ possible variants of a 200 amino acid protein, compared to “only” ~10^80^ protons in the visible universe [1]. The preponderance of possible proteins have never been realized over the course of evolutionary history and this rich space presents a large palette with which to discover novel proteins with “super-natural” or even non-natural function [2]. The vast size of this space also presents a challenge since exhaustive traversal by either experiment or computation is intractable, and the estimated probability of a randomly selected sequence being functional has been estimated to be as low as 1 in 10^77^ [3]. Moreover, the higher or more specialized functionality desired, the rarer such sequences become [4–6]. The rational design of protein sequences with programmed function requires models of the sequence-function (i.e., genotype-phenotype) relationship and a means to guide sampling from this distribution to generate plausible candidate sequences with the desired functionality for experimental synthesis and testing [4, 7].

The protein sequence-function relationship is a complex mesoscale phenomenon for which we lack good theoretical models. Accordingly, significant effort has been invested into the development of data-driven and empirical approaches. The famous sequence hypothesis of Crick and Watson stipulates that within a particular environment the protein amino acid sequence dictates the three-dimensional protein structure that, in turn, dictates the protein function [8–10]. Historically, the sequence-structure mapping has been adopted as a proxy for the sequence-function relationship, with the functional design task reduced to the engineering of a particular three-dimensional fold (e.g., optimization of an enzymatic active site, engineering of a binding cleft). In recent years, deep learning networks exploiting modern tools such as equivariance-inducing architectures and diffusion models have broken new paths in computational protein structure prediction with atomic-level accuracy [11, 12] and, very recently, programmability of desired three-dimensional structures [7]. These technological advances have also powered direct learning of the sequence-function relationship using approaches such as recurrent neural networks [13, 14], variational autoencoders [15–21], generative adversarial networks [22], reinforcement learning [23], and transformers [24–31]. A number of models have demonstrated remarkable performance in functional prediction tasks such as fluorescence, stability, and epistasis [24, 32].

A protein design model should possess four key characteristics: (i) *accurate* learning of the sequence-function relationship, (ii) *generative* design of sequences under this learned mapping, (iii)*fast* and *transferable* model training, and (iv) the capacity for *unsupervised* training over unlabeled sequence data and *semi-supervised* retraining over labeled data. The first and second properties requires a sufficiently expressive and powerful model to learn the correlated patterns of amino acid mutations responsible for mediating function and a means to conduct “inverse” design wherein the relationship may be used to produce sequences with predicted function. Early quantitative structure property relation (QSPR) / quantitative structure activity relation (QSAR) models tended to employ hand-crafted sequence featurizations and relatively simple pseudo-linear model forms that limit the expressive power of the learned relationship and can require significant user intervention and/or domain knowledge [1]. Moreover, these techniques are typically discriminative as opposed to generative, meaning that an auxiliary process such as Monte Carlo simulation or simulated annealing is required to sample sequences from the learned mapping [33, 34]. Modern deep learning networks can, in principle, learn amino acid correlations of all orders to expressively represent the “syntax” of the sequence-function mapping and particular architectures can be made inherently generative. The third property is important to enable efficient training of frequently large and expensive neural network models and amortization of expensive training via transferability to multiple protein families. The fourth property is particularly germane to protein engineering applications where the vast size of protein sequence space and time and labor costs of experimental assays mean that the volume of labeled data points (i.e., sequences annotated with experimental measurements of the desired function) tends to be eclipsed by unlabeled data (i.e., unannotated sequences). The initial round of a typical engineering campaign may begin with no labeled sequence data and the first task for the data-driven model is to prospectively identify promising sequences for experimental testing. Accordingly, it is desirable for a model to admit training in a fully unsupervised manner and subsequent updating in a semi-supervised fashion.

Based on these desired criteria, a number of deep learning architectures have been employed for data-driven protein design. Two approaches in particular have received substantial attention: variational autoencoders (VAEs) [15, 16, 20, 21] and transformer-based models [24–31, 35]. VAEs are deep generative models comprising two consecutive neural networks [36, 37]: an encoder compresses the high-dimensional sequence data into a low-dimensional latent space that is then passed to a decoder whose task is to reconstruct the input sequences as the output of the network. The VAE is trained by variational inference to minimize a loss function that ideally balances reconstruction accuracy and regularization of the latent space under a prior distribution [36–38]. The loss function can also be augmented with a semi-supervised prediction head that regresses the assayed function of (some fraction of) the training sequences that have been subjected to experimental measurements [39–41]. This functionality enables iterative semi-supervised re-training of the network after conducting experimental synthesis and testing of novel synthetic sequences to update the model with data from non-natural regions of sequence space and also sculpt and induce functional gradients in the learned latent space [20]. The latent space serves as an information bottleneck that furnishes a projection of the sequences onto a low-dimensional manifold spanned by a small number of correlated patterns of amino acid mutations (i.e., “design rules”) that underpin the sequences in the training data. Annotation of the latent space by phylogenetic or phenotypic properties frequently reveal dimensions within the latent space to be correlated with phylogenetic evolution and/or protein function [15, 20]. The interpretable low-dimensional embedding is extremely valuable in both understanding and interpreting the distribution of sequences in the training data and, crucially, in providing a smooth, continuous, low-dimensional embedding to guide interpolative or extrapolative generative decoding of novel sequences designed to optimize desired protein functions [15, 20]. Despite these attractive properties, VAE models frequently necessitate that sequences be provided as fixed length vectors, which requires sequences to be arranged within multiple sequence alignments (MSAs) [42]. Although MSAs can convey predictive advantages in structure prediction [24], they can be laborious to construct, introduce biases in the placement of gaps, require updating in the face of novel sequence data, and limit applications to homologous families – families of proteins with a common evolutionary origin – that admit an alignment, thereby frustrating the transferability of models between families or as generic models for arbitrary proteins. This limitation to homologous families and fixed-length data can be alleviated through the use of convolutional or recurrent layers to featurize protein families prior to dimensionality reduction into the latent space [17], although this can present challenges in learning long-range mutational correlations.

Transformers are a deep learning architecture using the attention mechanism to learn many-body and long-range correlated patterns by self-supervised training commonly employing a masked language modeling objective [43]. A primary motivation for the development of these models was to solve the vanishing gradient problem that plagues learning of long-range correlations within recurrent neural network architectures while preserving the possibility of operating on variable length sequence data and auto-regressive decoding. These architectures occasioned a new generation of large language models (LLMs) that excel at natural language processing (NLP) tasks [44] and underpin a number of protein language models (pLMs) for protein functional prediction and design [29, 30, 35]. The high-level structure of a typical transformer employs an encoder block comprising a stack of self-attention layers incorporating position-dependent encoding to learn the correlated patterns of amino acid mutations defining the “syntax” of the protein sequence data within a latent space embedding. The latent space embedding may then be used either as an expressive featurization for high-accuracy downstream functional prediction tasks or, similar to a VAE, as a conditioning input for a decoder stack to generatively decode synthetic protein sequences [24, 32, 45]. Transformers are not limited to fixed length data, thereby dispensing with the requirement for MSAs and enabling training of generic pLMs over millions or billions of (non-homologous) protein sequences residing in large public databases such as UniProt [46] and BFD [47, 48]. This opens up the possibility for model transferability wherein a generic model is applied to a particular protein engineering task possibly after some limited fine tuning [25]. Sequence generation can also be conditioned on control characters or partial sequences to guide synthetic protein design [27, 49]. The dimensionality of the fixed-length latent space of typical transformer-based pLMs is (*n* × *d*), where *n* ≈ 512 is a typical maximum sequence length that can be accommodated and *d* = 1024 is a typical per token latent space dimensionality. The exceedingly high latent dimensionality is an inherent facet of transformer models that sacrifices easy interpretability of phylogeny and functional patterns and frustrates conditional generative design of synthetic proteins within particular regions of the latent space guided by functional gradients. Training is also typically very expensive with typical pLMs containing billions of trainable parameters that make even fine tuning an expensive endeavor.

In this work, we introduce the Protein Transformer Variational AutoEncoder (ProT-VAE) as a model that blends the relative merits of VAEs and transformers to achieve all four of the desired criteria outlined above. The high-level architecture of the model can be conceived as sandwiching a VAE between the encoder and decoder stacks of a transformer pLM. This architecture is enabled by the capacity of the transformer to operate on variable length data but furnish fixed-length latent representations. These representations can be conceived as expressive featurizations of protein sequences that serve as the input to the VAE. Within this sandwich model, the interface between the exterior transformer and the interior VAE is mediated by stacks of 1×1 convolutions acting as dimensionality reduction layers that compress and decompress the transformer latent space representation for processing by the VAE. The transformer and compression/decompression modules are generic, non-family specific networks that are trained over millions or billions of protein sequence training examples. The VAE is a lightweight network that is trained anew for each particular homologous protein family that is the subject of a design or engineering task. Training of the inner VAE for each specific family is fast, and is the only non-transferable component of the architecture. The primary hypothesis underpinning the ProT-VAE network is that homologous protein families lie on low-dimensional manifolds of substantially lower dimensionality than that of the transformer latent space, and that this space can be compressed and decompressed within a VAE to furnish a low-dimensional latent space suitable for interpretation, annotation, and guided generative design. The convolutional layers perform partial compression/de-compression in a generic, non-family specific manner, and the family-specific VAE performs the remaining compression/decompression into and out of the family-specific latent space. Essentially, the ProT-VAE thereby combines the properties of transformers as generic, transferable, and powerful featurizers capable of learning long-range correlations and operating on variable length sequence data, with the capacity of VAEs to furnish low-dimensional latent embeddings to guide generative sequence design and which can be quickly and iteratively retrained in an unsupervised or semi-supervised fashion [45]. We construct and train the ProT-VAE model using the NVIDIA BioNeMo framework and demonstrate its capacity as a powerful, extensible, and lightweight model for data-driven protein design in retrospective computational prediction tasks and prospective design of novel functional proteins that are subjected to experimental synthesis and testing.

## 2 Results and Discussion

### 2.1 Prior Art

Deep networks integrating transformers and VAEs have been previously pioneered in the context of a number of application domains. Wang and Wan developed the Transformer-Based Conditioned Variational Autoencoder (T-CVAE) wherein a VAE was used to learn a distribution over story plots and serve as a conditioning variable for the transformer decoder in story completion tasks [50]. Jiang et al. developed the Transformed Variational AutoEncoder by combining the Music Transformer and Deep Music Analogy to develop a model capable of learning long-range dependencies within musical melodies, furnish interpretable embeddings via a disentangling conditional VAE, and a means to transfer melody and rhythm between contexts [51]. Li et al. and Park and Lee employed, respectively, pre-training and fine tuning approaches to mitigate posterior collapse in a transformer VAE model for text [52, 53]. Arroyo, Postels, and Tombari proposed the Variational Transformer Network (VTN) as a synthesis of self-attention encoders and decoders within a VAE architecture for layout detection and generation [54]. Henderson and Fehr employed VAEs as an information bottleneck regularizer for transformer embeddings and used this model to embed and generate text within a nonparametric space of mixture distributions [55]. Our work is most closely related to the recent work of Castro et al., who introduced the Regularized Latent Space Optimization (ReLSO) approach for data-driven protein engineering [40]. The jointly trained autoencoder (JT-AE) architecture underpinning this approach comprises a transformer encoder, low-dimensional projection into a latent space bottleneck, 1D convolutional neural network (CNN) decoder, and fully-connected network to predict function from the latent space embedding. The ProT-VAE introduced in the present work shares similarities with the ReLSO approach in the use of a transformer-based featurization and subsequent compression of this encoding into a low-dimensional latent space, but ProT-VAE is distinguished by its use of an attention-based decoder stack for sequence generation to efficiently learn long-range correlations, its use of a transferable encoder and decoder such that the lightweight VAE is the only model component that requires retraining for each protein engineering task, the capacity for unsupervised/semi-supervised training that does not require all sequences to have attendant experimental measurements, and demonstration of the technique in wet lab testing of synthetic protein sequences.

### 2.2 Architecture of ProT-VAE

Our proposed ProT-VAE model fundamentally behaves as a variational autoencoder that operates directly on unaligned sequences of a homologous protein family to yield a low-dimensional, generative, and organized latent space from which new proteins with desired properties can be generated. A schematic of the ProT-VAE architecture is given in Fig. 1. The architecture has three key components or blocks for encoding and three for decoding.

**Fig. 1.**
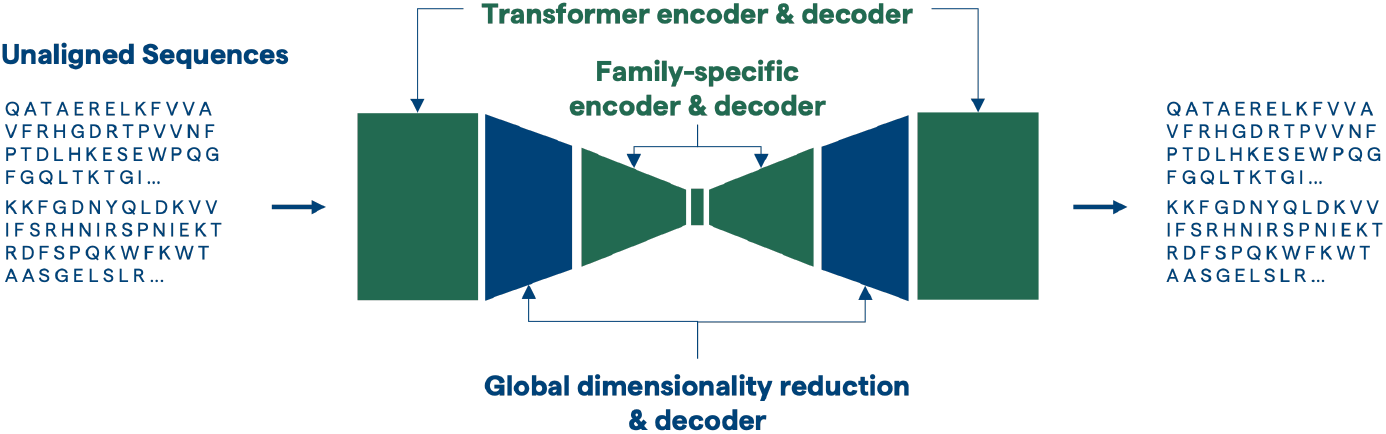
Overview of the ProT-VAE model architecture. A large language model encoder and decoder serve as the outermost blocks for a VAE. Unaligned protein sequences are first fed into a pLM encoder. The resulting large hidden states are efficiently compressed into a much more compact representation that still retains all of the information using a dimensionality reduction block composed mainly of a series of 1 × 1 convolutions. This reduced representation is finally fed into a series of fully connected layers ending in a low-dimensional bottleneck. The reduced dimensions are expanded the same way in reverse during the decoding process to reconstruct unaligned protein sequences. The pLM and convolutional compression/de-compression reduction layers are trained on large libraries comprising millions of protein sequences whereas the VAE is trained on specific families for each particular design task.

The first block is a pretrained transformer-based T5 encoder and decoder model referred to as ProtT5nv. ProtT5nv is readily available in the NVIDIA BioNeMo framework, currently available under early access and planned for future open source release [56, 57]. The model has 12 layers, 12 attention heads, a hidden dimension of 768, and 198M parameters. Pre-LN layer normalization and GeLU activation are used throughout the model. Additionally, encoder embeddings and decoder projections to logits are shared in this architecture. The model was trained with a maximum input sequence length of 512 and a masking probability of 15%. Unsupervised mask prediction was used as a training objective. Dropout was set to 0.1 during training.

ProtT5nv was trained starting from a T5 model pretrained on NLP data. Parameters of all layers from the NLP-pretrained T5 model, except for encoder and decoder embeddings, were used to initialize the ProtT5nv models weights. The original NLP-pretrained T5 model had a dictionary of 29,184 tokens, while the ProtT5nv model only required 128 tokens, including 96 sentinel tokens. ProtT5nv encoder embeddings were therefore initialized with 128 first encoder embedding vectors from the NLP-pre-trained T5 model. Then, decoder projections to logits were tied to encoder embeddings.

After initialization, the model was further trained with protein sequences from UniRef50, release 05/2022 [58]. Protein sequences longer than 512 amino acids were removed, resulting in approximately 46M samples. The sequences were randomly split with 4.35K in validation, 875K in test, and the remaining in train. ProtT5nv model was trained using data parallelism on 224 V100 GPUs for 58 epochs (approximately 1M iterations) using a micro-batch size of 12 protein sequences per GPU. Inverse square root annealing was used as a learning rate scheduler, with a minimum learning rate of 0.0 and 10000 warmup steps. Fused Adam optimization was used with parameters *β*_1_ = 0.9, *β*_2_ = 0.999, and weight decay=0.01.

The second block is a generic dimensionality reduction block that serves to efficiently compress the ~300,000-dimensional transformer hidden state into a more parsimonious intermediate-level representation. These intermediate layers are also pretrained on large protein databases and do not require significant changes per protein family. For this work, these layers were pretrained using a mean squared error (MSE) reconstruction objective on Uniprot. This block consists of several stacks of dimensionality reduction layers, where each layer comprises (i) 1×1 convolutions, (ii) LayerNorm, (iii) GeLU activations, and the filter size is incremented at each step. In the results presented here, three layers were used with filter sizes of 512, 256, and 64 in the encoding side and 256, 512, and 768 in the decoding side. These layers serve as a generic dimensionality reduction which are very parameter efficient, fast to train, and do not suffer from significant loss in information content. When trained and validated on large datasets of proteins, reductions of 16× or more are possible without any noticeable degradation of reconstruction quality. In the current implementation, the full output of the transformer hidden state, including the positions corresponding to padding tokens, are compressed to create a resultant 32,768-dimensional intermediate representation serves as a much more compact starting point for the family-specific VAE layers to operate upon and allows for better latent space generation with fewer total network parameters.

The third block is a three layer fully-connected maximum mean discrepancy variational autoencoder (MMD-VAE) employing ReLU activations [59] that takes the flattened output of the dimensionality reduction block and compresses it into a protein family-specific, low-dimensional latent space. Due to its specificity, this network is initialized and trained from scratch for each target protein family of interest for a particular design task and furnishes the functionally organized, low-dimensional, generative manifold from which synthetic proteins are designed.

### 2.3 Training of ProT-VAE

A pretrained ProtT5 was obtained from the previously described BioNeMo checkpoint. The intermediate dimensionality reduction layers were pretrained on Uniprot with using Adam optimizer with a learning rate of 0.0001 on an MSE objective on reconstruction of the full T5 hidden state. T5 layers were frozen during this pretraining. Family-specific VAE layers were trained on full, unaligned sequences belonging to a single homologous protein family. Sequences were fed in with 15% of positions randomly masked and 20% of masks corrupted to different amino acids. The training objective was the reconstruction of the original sequence, optimized using Adam with a learning rate of 0.0001.

### 2.4 Latent Representation and Organization

We demonstrate and test ProT-VAE in applications to two protein families: the Src homology 3 (SH3) protein family involved in diverse signaling functions within cells, and the phenylalanine hydroxylase (PAH) enzyme that catalyzes conversion of phenylalanine to tyrosine. We test the capability of the ProT-VAE model to learn meaningful and interpretable latent spaces organizing protein sequences by ancestry and function, to make accurate predictions of protein function from the learned latent space, and to generatively design novel synthetic sequences with function commensurate or superior to natural sequences and with high sequence divergences from the natural training data.

#### 2.4.1 SH3

The Src homology 3 (SH3) is a family of small beta folds that mediate protein signaling within cells by binding to type II poly-proline peptides with sequences N-R/KXXPXXP-C or N-XPXXPXR/K-C [60, 61]. SH3 domains have evolved to perform a variety of functions within various organisms by evolving differential binding specificities, resulting in a number of distinct paralogs (i.e., homologous proteins performing different functions within the same species) within the SH3 family. Recent work by Lian et al. trained VAEs over an MSA of ~--5,300 SH3 homologs to develop a deep generative model for synthetic SH3 design [20]. The VAE learned an unsupervised three-dimensional latent space embedding in which the natural sequences demonstrated an emergent hierarchical clustering by phylogeny and function. The Sho1^SH3^ domain in *Saccharomyces cerevisiae* (i.e., baker’s yeast) mediates transduction of an osmotic stress signal by binding a Pbs2 ligand that activates a homeostatic response to balance the osmotic pressure by intracellular production of glycerol [62]. A high-throughput *in vivo* osmosensing assay was developed to measure the relative enrichment of deep sequencing counts of *S. cerevisiae* Sho1^SH3^ knock-outs into which mutant SH3 genes designed by the VAE were transformed. The normalized relative enrichment (r.e.) is shown to quantitatively report on the binding free energy of the mutant SH3 with the Pbs2 ligand. The assay demonstrated that natural Sho1^SH3^ orthologs reside within a localized cluster within the VAE latent space and generative design of mutant sequences in the vicinity of this cluster conferred equal or superior high osmolarity protection to wild type Sho1^SH3^. As a first demonstration of the ProT-VAE model we test its capacity to learn interpretable latent space embeddings of the SH3 family organized by phylogeny and function without the need for an MSA. The inference of phylogenetic and functional relationships within a learned latent space is a pre-requisite to subsequent data-driven functional protein design.

#### 2.4.2 Functionality

We fine-tune the inner two VAE layers of the ProT-VAE model on the SH3 dataset reported in Ref. [20] employing a 6D latent space. The results are displayed in Figure 2. The latent space is plotted in 2D projections in three panels, colored by sequences with high r.e. scores. The cutoff based on the dataset is sequences with a normalized r.e. > 0.6. Based on the colored projections in Figure 2A, the high activity sequences are clustered in all dimensions. We featurize the activity into a binary dataset of high and low activity. Using the binary label, we evaluate the generative potential of the model. Due to the low number of high activity sequences in the dataset as can be seen from Figure 2B, the model has a low average reconstruction of the validation sequences. Using the latent space embedding and the binary labels, a logistic regression was trained via 5-fold cross validation. The results of the regression are shown in the receiver operating characteristic (ROC) curve in Figure 2C. The ProT-VAE slightly outperforms the MSA-based VAE model in predicting functional performance with an AUC of 0.98 compared to 0.95. This result validates the capacity of the ProT-VAE model to accurately predict protein function without the requirement for an MSA.

**Fig. 2.**
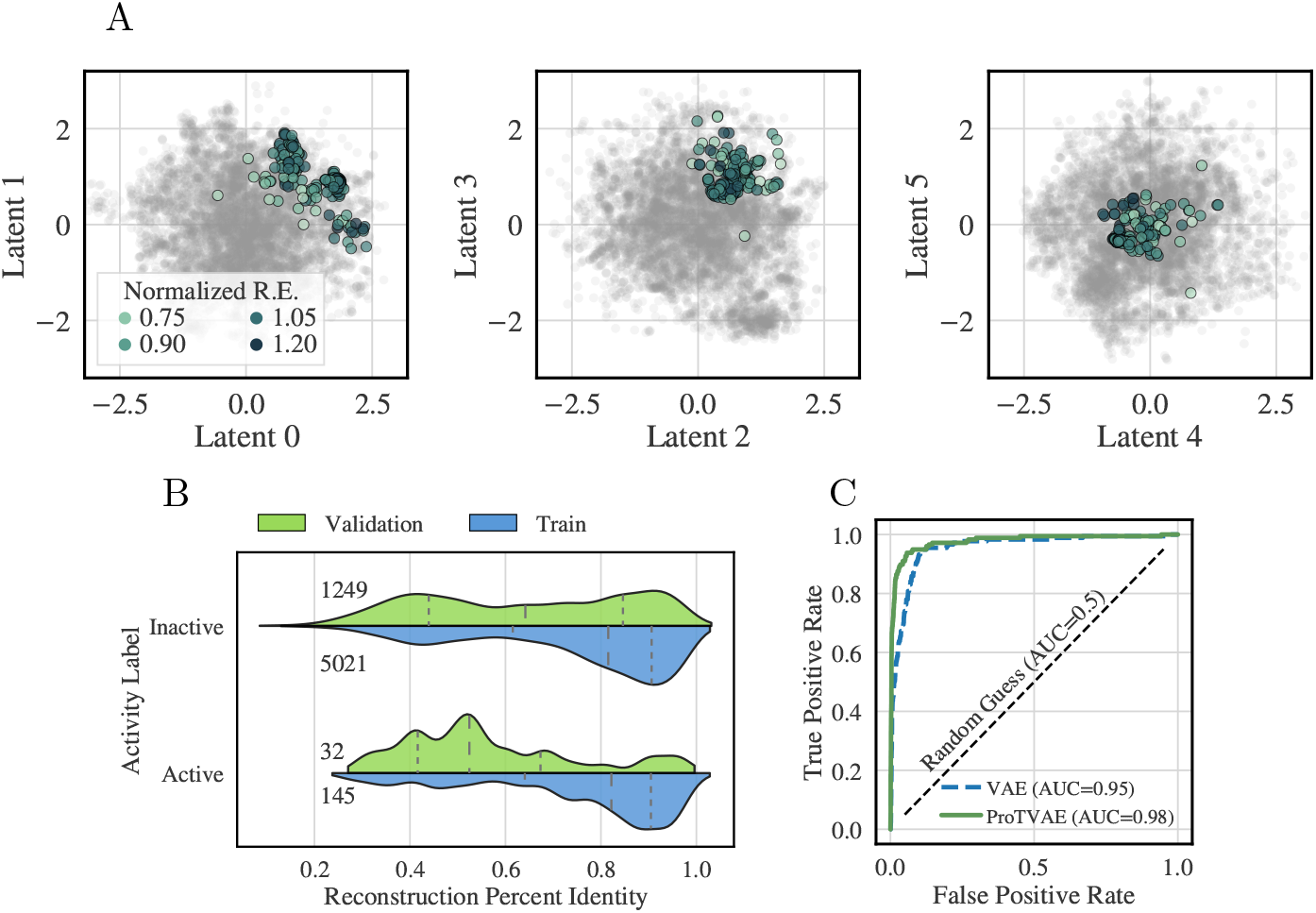
ProT-VAE organizes the SH3 family by functional activity without an MSA. A) Two-dimensional projections of the latent space are shown colored by normalized relative enrichment (r.e.), where darker points correspond to more active sequences. Note the clustering of these sequences in all three projections. B) Reconstruction strength of SH3 sequences characterized by activity. The distributions for each classification are represented by violin plots separated into those in the training set (bottom-half in blue) and those in the validation set (top-half in green). The number of sequences within each classification and dataset split are printed on the left of the plot. Dashed lines correspond to the 25%, median, and 75% quartile ranges moving from left to right. C) Prediction of functionality via latent space trained classification model. A logistic regression classifier was trained via 5-fold cross validation, using the latent space coordinates and the binary activity labels. The resultant ROC of the classifier is shown, demonstrating that the latent space is organized such that protein functionality is localized within the latent space. The green solid line corresponds to the ProT-VAE model, the blue dashed line corresponds to the previous MSA-based VAE model, and the black dashed line is the null hypothesis. In the legend, we show the respective AUC scores of each of the classifiers.

#### 2.4.3 Phylogeny

In addition to the activity benchmark, we seek to reproduce the phylogenetic separation within the latent space previously reported in the alignment-based VAE model [20]. At first glance, there was shown to be no correlation between phylogeny and latent space encoding, however, this was actually shown to be the result of a hierarchical organization: the model separates by paralog group, and then phylogeny is separated within each paralog cluster. We seek to reproduce the same results with ProT-VAE to show the parity between MSA-based and MSA-free methods (Figure 3). In Figure 3A, we demonstrate that the ProT-VAE model is also unable to visually separate phylogeny between Asomycota and Basidiomycota in any latent dimension. Given this result, we seek to establish the same hierarchical effect as shown in the MSA-based model. In Figure 3B, we show that there is clear qualitative clustering of the paralog groups of the SH3 family between Abp1, Rvs167, Sho1, and Bzz1. From here, we examine the capability of the model to separate phylogeny within each paralog. In Figure 3C, we separate the Sho1 paralogs by phylogeny, now noting a key separation, especially in the first two latent dimensions, just as has been shown previously. We show the ability of the model to cluster other annotated paralog groups by phylogeny in Figure A1. The capacity of ProT-VAE to learn functional and phylogenetic separation within the latent space without the need for MSAs demonstrates its capacity to learn the correlated patterns of amino acid mutations underpinning the ancestral history and functional performance of sequence ensembles, and opens the door to alignment-free generative design and engineering of novel synthetic proteins. We now assess the performance of the model in these tasks in the context of design of a therapeutic protein.

**Fig. 3.**
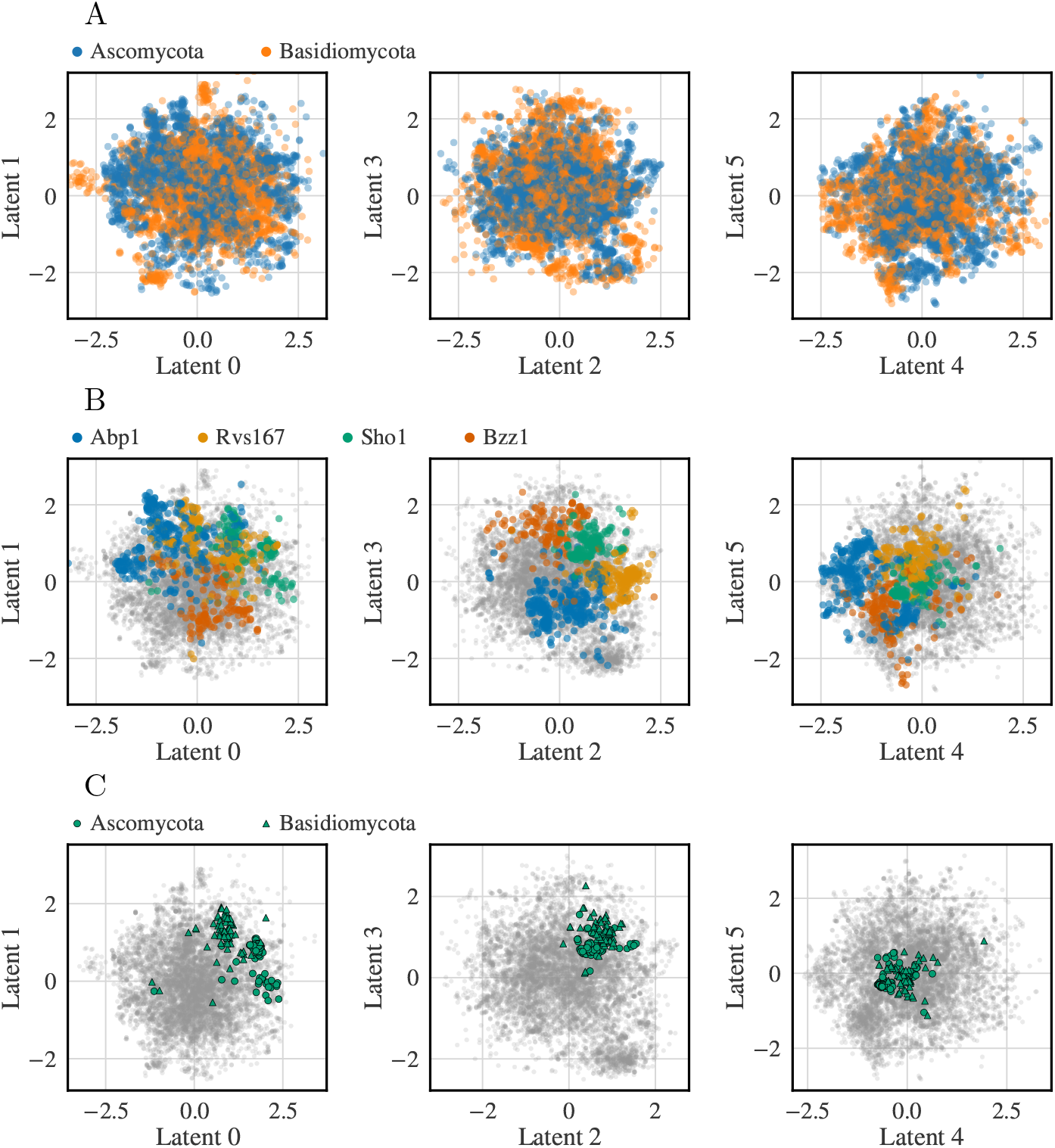
ProT-VAE hierarchically organizes SH3 family first by paralog group and then by phylogeny. A) Two-dimensional projections of the latent space are shown colored by two phylogenetic groups, Ascomycota and Basidomycota with no apparent organization. B) Two-dimensional projections of the latent space colored by paralog groups (Abp1 in blue, Rvs167 in orange, Sho1 in green and Bzz1 in yellow) exhibit strong clustering. C) Considering just the Sho1 paralog group, we then color by phylogeny again and observe good separation of Ascomycota and Basidomycota within this paralog group. Analogous plots for the other paralog groups are presented in Figure A1.

### 2.5 PAH

To further evaluate the ProT-VAE model, we examine a second protein family with therapeutic properties. In particular, we consider the family of aromatic amino acid hydroxylases (AAAH), specifically phenylalanine hydroxylase (PAH). Human PAH (hPAH) is an enzyme that catalyzes the catabolism of one amino acid, phenylalanine, into another, tyrosine, by hydroxylation of the Phe side chain [63]. This reaction is critical in eliminating surplus phenylalanine and producing tyrosine as an essential precursor for the production of hormones, neurotransmitters, and pigments. Starting from a human PAH variant, 2PAH [64], a psiBLAST was run to find a dataset of homologous proteins, resulting in a dataset of 20,000 sequences. Using annotations from the non-redundant protein database, the dataset was charactized both by sub-strate specificity and phylogeny. To effectively test the encoding strength of the ProT-VAE, we fine-tuned the inner two VAE layers of ProT-VAE under an 80-20 training-validation split. We project all sequences into a 6D latent space for evaluation.

#### 2.5.1 Substrate Specificity

Our first task for the model is testing its encoding strength with respect to organizing the substrate specificity annotations of the sequences. In Figure 4A, the latent space is shown in two-dimensional projections, colored by substrate specificity. From the figure, we observe good separation and clustering of the most labeled functional substrates of the AAAH family: tryptophan, phenylalanine, tyrosine, and henna. To test the generative aspect of the model, we look at the percent identity of reconstructed sequences. To evaluate this, we pass sequences through the encoder of the model and then decode from the latent space, calculating the percent identity between the initial sequence and the decoded sequence. We show the distributions of reconstruction by data split and substrate in Figure 4B, along with associated counts in each split. In these distributions, we note that there is a strong parity between the training set and validation set in reconstruction strength. For example, henna hydroxylation enzymes consist of the smallest amount of proteins (~ 1%) yet the validation fraction reconstruction is comparable to the training set median and both are above 90%. Using the latent space encoding, we trained a k-nearest neighbors classifier with k=5 and five-fold cross-validation. In Figure 4C, we show the quantitative separation of the latent space embedding via a confusion matrix of the classifier trained on the embeddings. Despite the unbalanced class labels, the model is able to both generate these sequences with high reconstruction accuracy. In addition, we note that with unsupervised learning, the model is also able to predict the substrate specificity with high accuracy based on the learned functional organization of the latent space.

**Fig. 4.**
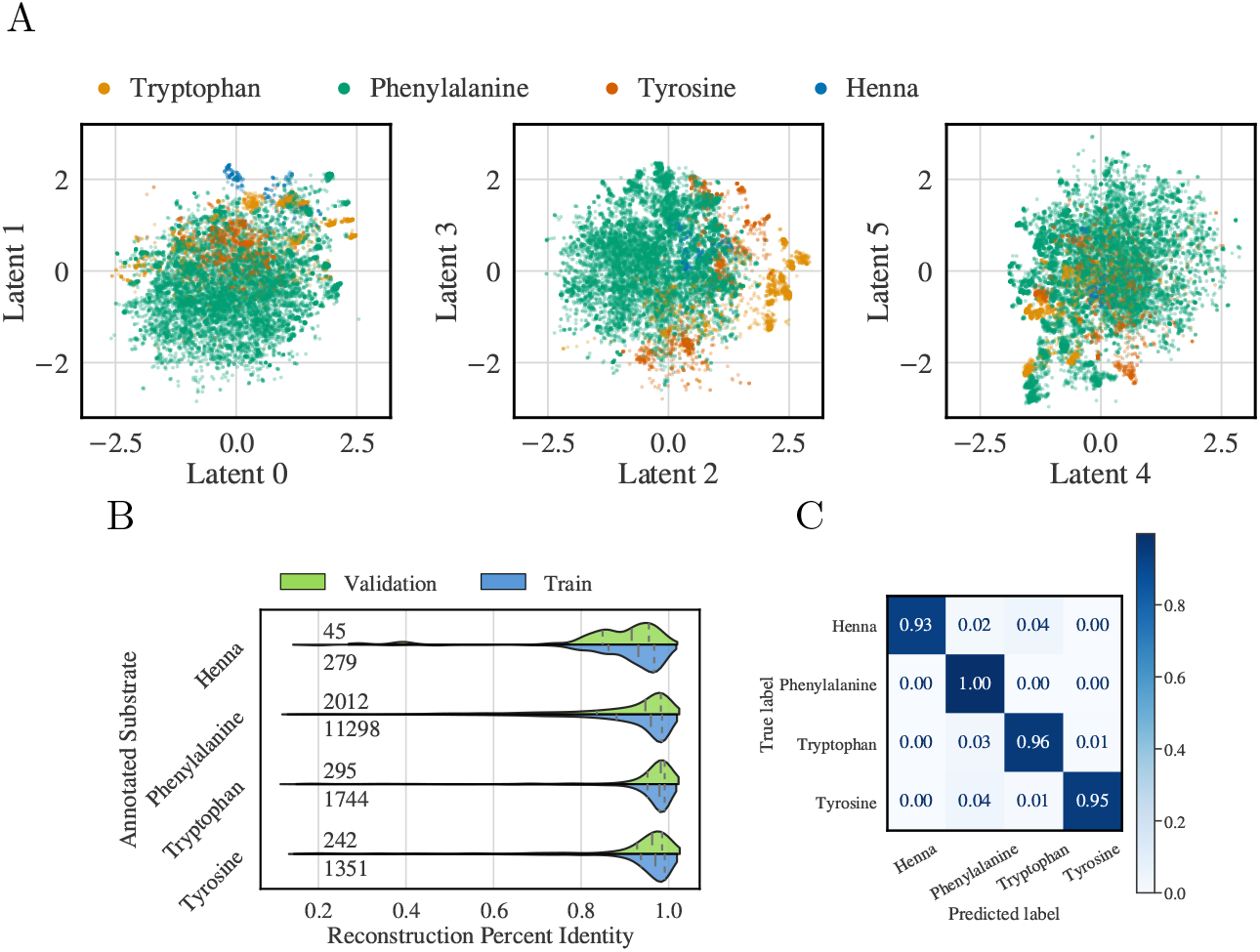
ProT-VAE organizes the AAAH family by function without needing an alignment. A) Two-dimensional projections of the latent space are shown colored by substrate specificity. B) Reconstruction of AAAH family sequences characterized by substrate specificity. The distributions for each classification are represented by violin plots separated into those in the training set (blue) and validation set (green). The number of sequences within each classification and dataset split are printed on the left of the plot. Dashed lines correspond to the 25%, median, and 75% quartile ranges moving from left to right. C) Prediction of functionality via latent space trained classification model. A k=5-nearest-neighbors classifier was trained via five-fold cross validation, using the latent space coordinates. The resultant confusion matrix of the model is shown, demonstrating that the latent space is organized such that protein functionality is localized.

#### 2.5.2 Phylogeny

While prediction functionality of the PAH protein is of importance, there are a myriad of phenotypes associated with each protein in the dataset. A key task in the design of therapeutics, for example, is reduction of immunogenic responses. In this vein, it is necessary to investigate the predictive capability of the model over multiple tasks, much in the same way that language models are evaluated on general benchmarks such as GLUE. [65] We break down the AAAH dataset by phylogeny at the phylum level and, using the same latent space, annotate the embedding by phylogeny. The results are displayed in Figure 5. Similar to the substrate specificity results, we observe good clustering the of the top phylum labels (Figure 5A). In addition, we see high reconstruction accuracy of all classes, even though there is a substantial imbalance across the class labels (Figure 5B). Such a result is encouraging for the generative potential of ProT-VAE for the *de novo* design of proteins with specific functionality for a specific host. This is exceptionally useful in, for example, the potential humanization of proteins for therapeutics with specific activity or specificity. We also note that by training another k=5-nearest neighbors classifier on the phylum labels with five-fold cross validation, we achieve high prediction accuracy (Figure 5C).

**Fig. 5.**
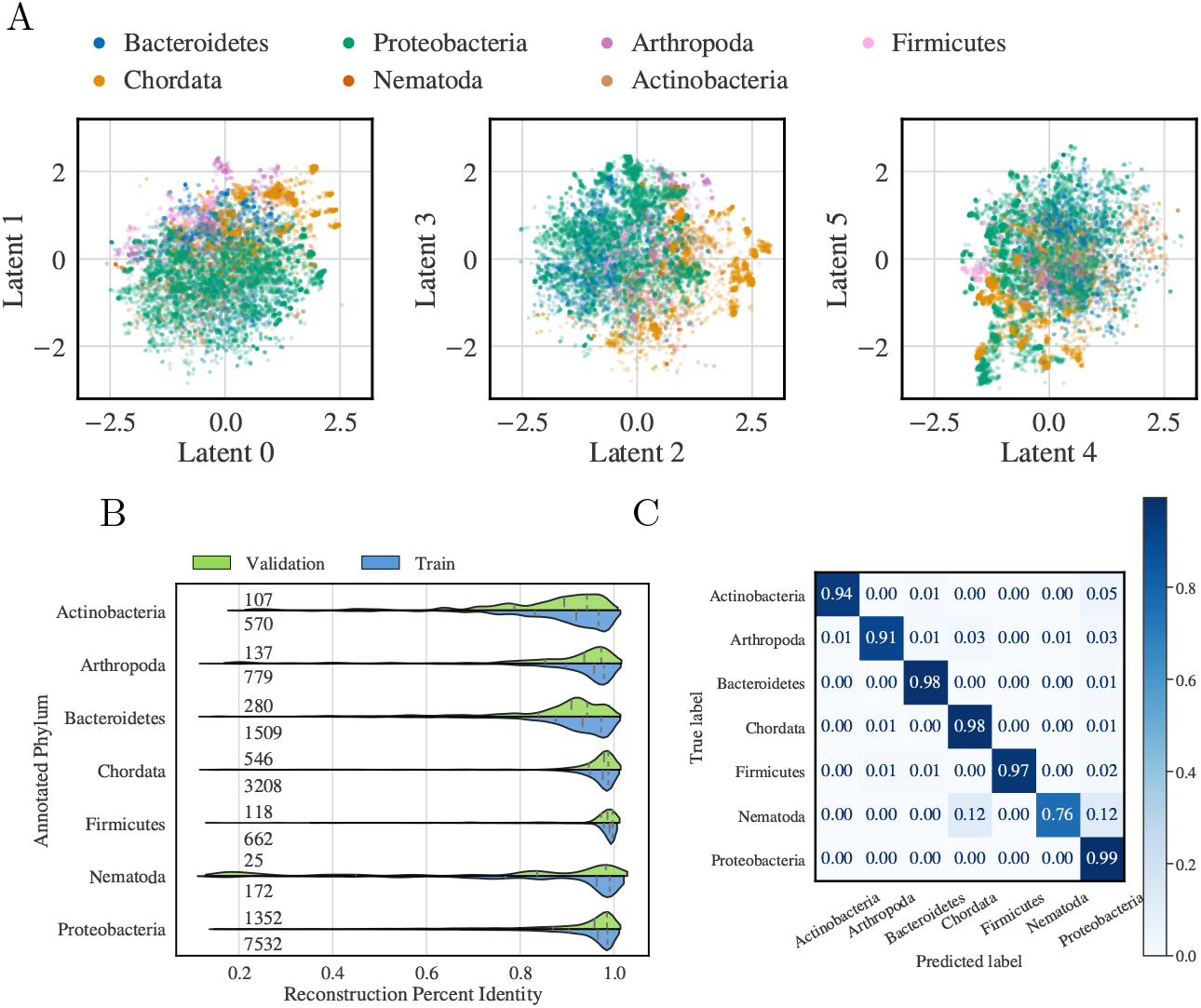
ProT-VAE organizes the AAAH family by phylogeny without needing an alignment. A) Two-dimensional projections of the latent space are shown colored by phylum characterization. B) Reconstruction of AAAH family sequences characterized by phylum. The distributions for each classification are represented by violin plots separated into those in the training set (blue) and validation set (green). The number of sequences within each classification and dataset split are printed in the left of the plot. Dashed lines correspond to the 25%, median, and 75% quartile ranges moving from left to right. C) Prediction of phylogeny via latent space trained classification model. A k=5-nearest-neighbors classifier was trained via five-fold cross validation, using the latent space coordinates. The resultant confusion matrix of the model is shown, demonstrating that the latent space is locally organized such that protein phylogeny is localized in the learned latent space.

#### 2.5.3 Latent Space Interpolation

A hallmark of a smooth latent space suitable for optimization and generative design is sensible interpolations. To test the smoothness of these latent spaces we performed two separate interpolations: one between similar phylogeny but different substrates, and one between different phylogenies acting on the same substrate. The substrate path was traversed between the 2PAH human PAH (hPAH) and a human tyrosine hydroxylase (hTyrH), while the phylogeny path was between the same hPAH and a flavobacteriacaea PAH sequence (bacPAH). We interpolate 50 points with spherical linear interpolation (SLERP) [45] and show the results of both interpolations in Figure 6.

**Fig. 6.**
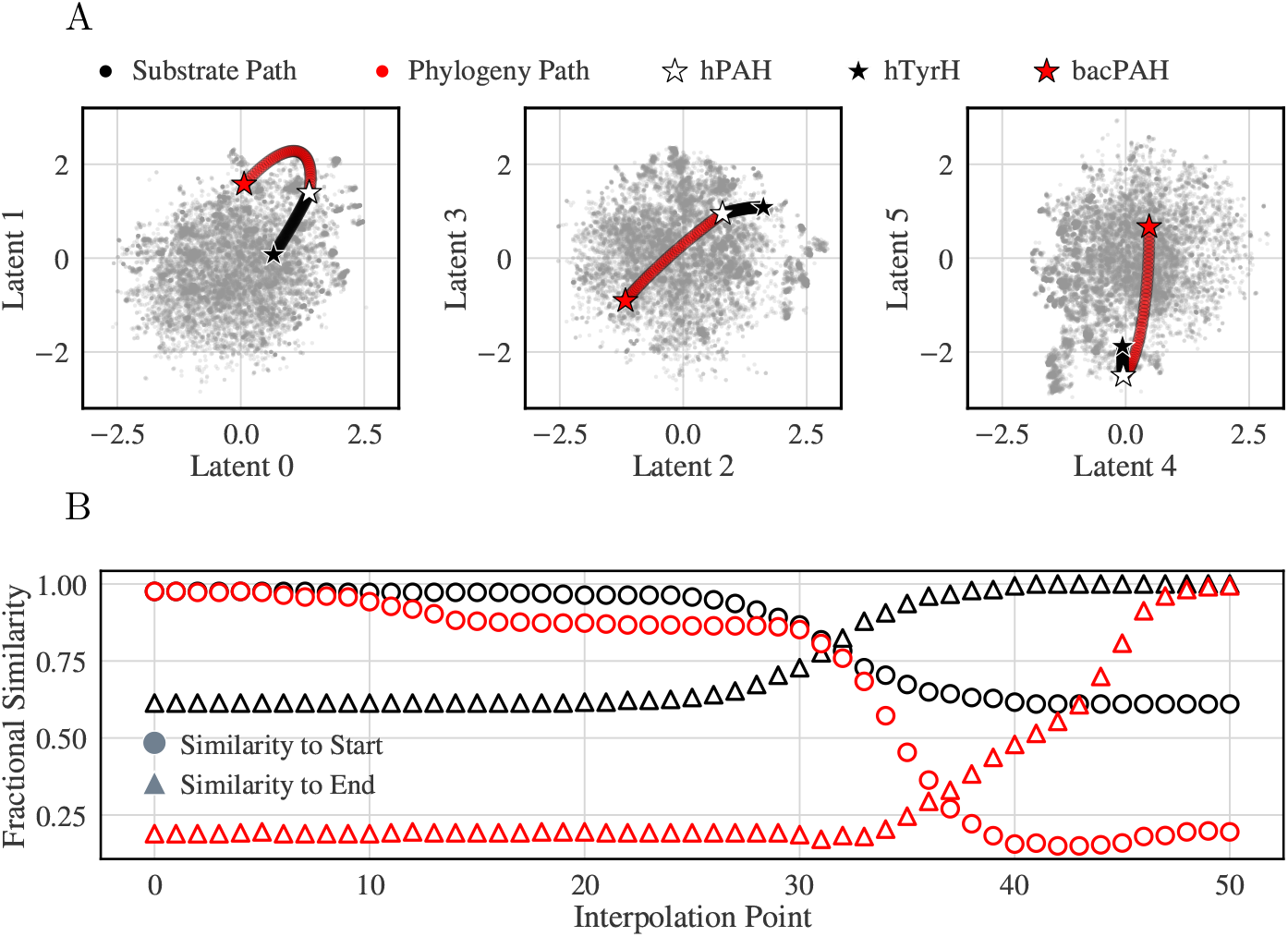
Traversal in latent space smoothly transitions between sequences. A) Two-dimensional projections of the spherical interpolation in the latent space. There are two paths represented: the phylogeny path, in red, which traverses between a human PAH (hPAH) represented by a white star and a bacterial PAH (bacPAH) represented by a red star, and a substrate specificy path, in black, which traverses between the same hPAH and a human tyrosine hydroxylase (hTyrH) B) A graph showing the steady transition from the first sequence into the second as the interpolation proceeds for both paths. Circles represent the percent identity of the sequences in the path with reference to the start and triangles represent similarity to the end sequence.

In Figure 6A, we visualize both interpolations in the latent space, colored by interpolation path. Interestingly, there is a longer traversal path for the phylogeny interpolation than for substrates, which is also correlated with larger sequence similarity changes. At each point, we reconstruct the sequence through the decoder to evaluate the sequence similarity of these novel sequences (Figure 6B). Despite unique paths of different lengths, there is a smooth transition within both paths that exhibits no sharp transitions. It is also of note that the phylogeny interpolation covers a range of 85% difference in sequence, and yet still results in a smooth interpolation. With this result we are encouraged not just by the organization of the ProT-VAE model to organize information with its embedding, but also to generate a smooth representation of unique sequences for generative design and experimental testing.

#### 2.5.4 Protein Design

Given the strong predictive performance of the model, we now proceed to evaluate its potential for *de novo* sequence design. Specifically, we test the capabilities of ProT-VAE to furnish interpretable and generative latent spaces by performing generative design of synthetic human-like PAH proteins for experimental evaluation with a plate-based assay. Given the strong predictive performance of the model, we now proceed to evaluate its potential for *de novo* sequence design. Specifically, we test the capabilities of ProT-VAE to furnish interpretable and generative latent spaces by performing generative design of synthetic human-like PAH proteins for experimental evaluation with a plate-based assay. To generate these sequences, we chose a conservative approach of local sampling around hPAH in latent space by fitting a multi-dimenisional Gaussian centered around hPAH. We filtered out sequences that were duplicates of natural sequences and imposed a maximum similarity of any generated sequence to any natural sequence in the training data of 99%. To localize the generated sequences in both latent space and sequence space, we also placed a cap on the maximum number of mutations (i.e., substitutions, insertions, deletions) in the generated sequences away from hPAH of 140 of the 333 wild-type positions corresponding to a minimum sequence similarity of 58%. Under these criteria, 190 sequences containing a maximum of 133 mutations away from hPAH and spanning a range of lengths of 223-339 residues were chosen for experimental synthesis and testing.

The assay is designed in 96-well plates using a Biotek plate reader to evaluate the fluorescence of tyrosine produced over a defined time period. The efficiency of catalytic conversion is calculate as the maximum velocity of the reaction, normalized by enzyme concentration in the well. The maximum velocity is calculated by taking the derivative of the curve and finding where it is at a maximum. These values are then normalized by the average hPAH velocity and converted into a fold over wild-type (FOWT) measurement, for easy comparison. Full details of the assay are provided in the Methods. We show the activities of the designed sequences in Figure 7.

**Fig. 7.**
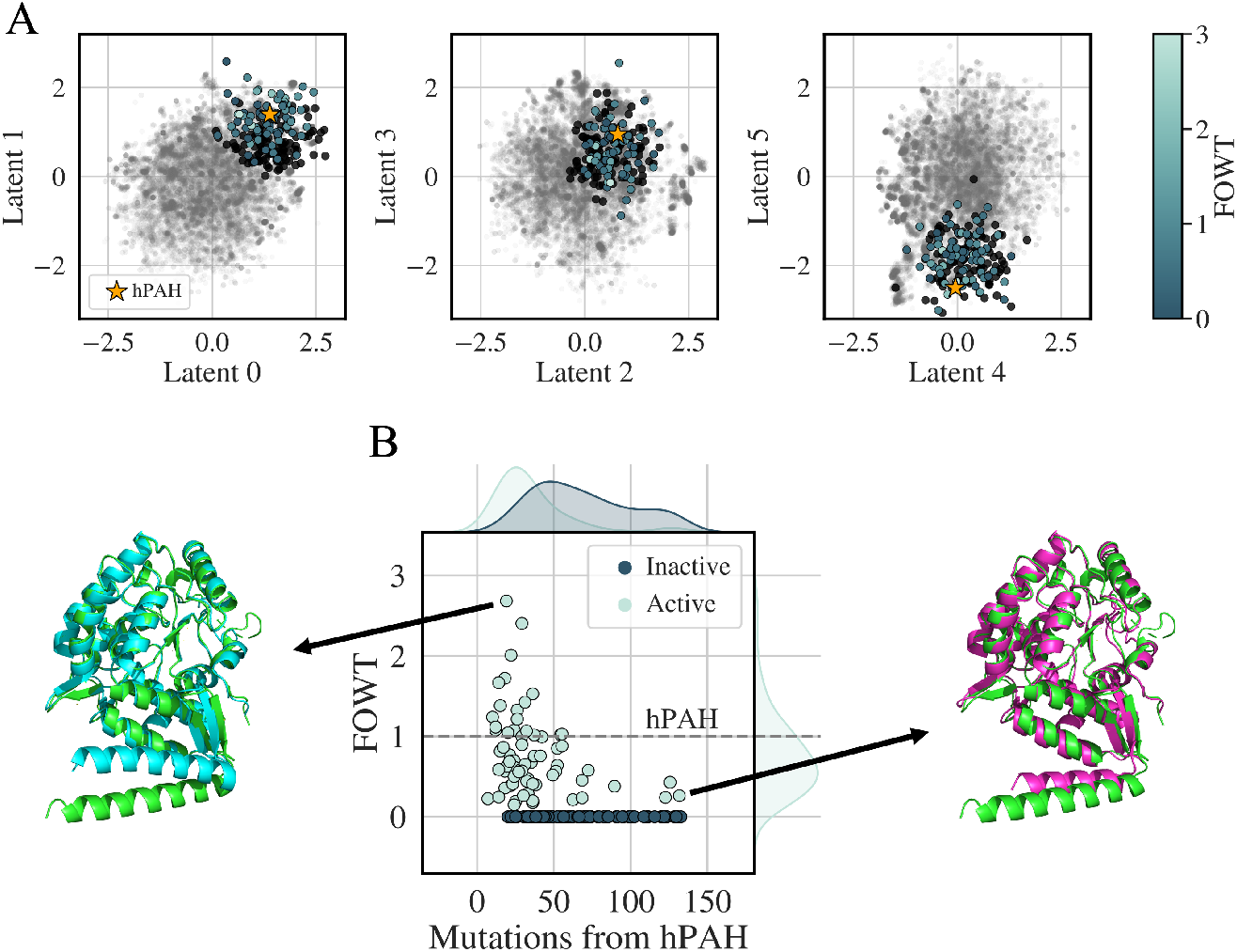
Generative design of ProT-VAE. There were 190 proteins locally designed around human PAH (hPAH) for experimental testing. A) The latent space organized by activity in fold-over-wild-type (FOWT). Color corresponds to activity; sequences in black are inactive. hPAH is indicated by a gold star. The training data for the model is shown in gray. B) Activity distribution of the designed sequences as a function of similarity to hPAH. We show similarity as number of mutations. We present the predicted structures generated by AlphaFold of the highest-activity synthetic mutant (teal), and the most highly mutated functional synthetic mutant (purple), each aligned with the natural hPAH (green).

In Figure 7A, we show the sequences annotated by their relative activity, FOWT, projected into the latent space. The gray points represent the training data to show the overall latent space projections. In black, we show the inactive sequences, which occupy the same regions of space as the active. All other sequences that are active are shown according to the colorbar to the right of the projections. From the figure, we note that the ProT-VAE model is capable of MSA-free *de novo* design of synthetic PAH proteins from a low-dimensional generative and interpretable latent space. Of the proteins assayed, 69 showed activity and 19 were more active than hPAH, with a maximum activity increase of 2.5× that of hPAH. In Figure 7B, we show the activities of the generated sequences as a function of the number of mutations from hPAH. Additionally, we predicted structures for the highest active variant and the most mutated variant that showed activity using AlphaFold [11]. The structural models suggest that the synthetic mutants preserve the native fold of the wild-type natural hPAH despite the ProT-VAE model being furnished no structural information. While the majority of the active sequences are very similar (< 30 mutations), we do note that the ProT-VAE model is capable of generating highly mutated sequences (> 100 mutations, highest active at 130 mutations) that are still functional. This dynamic range of generative design is highly non-trivial from a protein engineering standpoint and shows the engineering power that ProT-VAE provides. We are encouraged by the ProT-VAE model, not just for its embedding strength, but the ability of the model to generate novel, highly active, and diverse sequences.

## 3 Conclusions

In this work we introduce ProT-VAE, an accurate, generative, fast, and transferable model of the sequence-function relationship for data-driven protein engineering. By blending the desirable features of transformers and VAEs, the model admits alignment-free training in an unsupervised or semi-supervised fashion and furnishes interpretable low-dimensional latent spaces that facilitate understanding and generative design of functional synthetic sequences. The ProT-VAE model comprises a VAE to distill task-specific information from generally pretrained, attention-based transformer encoder and decoder stacks with the aid of intermediate compression/decompression blocks. Pre-trained transformer encoder and decoder stacks are taken from the 198 million parameter ProtT5 model trained over approximately 46M unique protein sequences within the UniRef50 (release 05/2022) database after clustering, truncation, and splitting. This model is made available within the NVIDIA BioNeMo framework [56, 57] and will be open-sourced in future releases. The model is, however, not dependent on the particular choice of encoder and decoder. We perform in-house training of generic compression/decompression blocks to map back and forth between the ~300,000-dimensional ProtT5 latent space and a 32,768-dimensional compressed representation that serves a fixed-length input to a task-specific VAE. The transformer and compression/decompression stacks are transferable and generic models that need only be trained once over large libraries of diverse protein sequences and can be conceived of as furnishing expressive fixed-length featurizations of arbitrary proteins from unaligned sequences. Only the lightweight VAE requires training anew for each protein engineering task and furnishes a smooth, low-dimensional latent space that furnishes interpretable understanding and a springboard for conditional generation of synthetic protein sequences with engineered function.

We validate ProT-VAE in applications to two different protein families: SH3 and PAH. We show that latent space organization is an excellent predictor of both phylogeny and function in both systems, allowing design of proteins with desired substrate specificity. We then show that sequences smoothly transition in latent space in interpolations performed on PAH. Finally, we design and experimentally validate new PAH sequences with more than 100 mutations and up to 2.5× increase in activity over the human PAH.

These tasks show that the learned latent space in the bottleneck layer at the heart of the network naturally organizes protein sequences according to ancestral history and biological function, exposes functional gradients in annotated sequences, supports high-accuracy regression models to predict the function of novel sequences, and enables the design of synthetic sequences by interpolative decoding and conditional generation along latent space pathways between natural sequences. Novel proteins generated by ProT-VAE exhibit substantial sequence diversity from any natural sequences in the training data yet possess high functionality as verified by experimental synthesis and functional assays.

ProT-VAE represents a powerful new architecture for data-driven protein engineering that can be deployed to generate synthetic proteins possessing high novelty and functionality for arbitrary design tasks. The only requirement is the availability of sufficient training data – typically ensembles of natural protein sequences – for stable training of a lightweight task-specific VAE to furnish a robust latent space embedding. Importantly, the model does not rely on the construction of multiple sequence alignments by using robust and expressive attention-based featurizations to eliminate the time, labor, and bias associated with alignment construction and efficiently learn of long-range mutational correlations. We have demonstrated ProT-VAE in downstream functional prediction and data-driven design of novel functional sequences after training the model on natural libraries. The model is, however, immediately deployable within subsequent rounds of a machine learning-guided directed evolution (MLDE) campaign by semi-supervised retraining of the VAE blocks on the synthetic sequences and their attendant functional assays [1, 4, 33, 35, 45, 66–71]. By virtue of the NVIDIA BioNeMo framework upon which the model is constructed, we can take advantage of pre-trained transformer encoder and decoder stacks and perform efficient retraining of the VAE for typical protein engineering tasks in just 12-24 GPU-h on commodity NVIDIA A100 GPUs, although inference and even retraining of the VAE can be performed on GPUs with as little as 10 GB of memory.

We envisage exciting applications of the ProT-VAE model as a powerful tool for a panoply of protein engineering and design applications. The transformer/VAE hybrid architecture is extremely powerful and generically extensible. We can conceive of numerous potential innovations on this architecture by incorporating alternative transformer encoders/decoders and different VAE loss functions and training protocols. In particular, we anticipate that the model may be straightforwardly applied to identification, conditional generation, or design tasks in diverse fields simply by switching out the encoder/decoder blocks for transformer models pre-trained over, for example, nucleic acids, synthetic polymers, small molecules, text, speech, music, or other sequence-based data that reside on low-dimensional manifolds.

## 4 Methods

### 4.1 Experimental measurement of PAH activity

#### 4.1.1 Methods

The insert was ordered from a gene fragment vendor. The 10× T4 DNA Ligase Buffer, Bsal-HF v2, and T4 DNA ligase were purchased from New England Biolabs. The molecular grade water was purchased from Corning. Mix & Go *E. Coli* Transformation Kit and Buffer Set was purchased from Zymo research. Aera Seal was purchased from Excel scientific. LB Broth was purchased from Gibco, and S.O.C. medium from Invitrogen, and 10x PBS was purchased from Gibco, Solulyse from AMSBIO, Lysonase from Novagen.

#### 4.1.2 High-Throughput (HT) Preparation and Screening

Highly competent BL21(DE3) cells expressing chaperone pGro7 (BL21(DE3)-pGRO7) were prepared in-house using a kit from Zymo Research (kit # T3002). Briefly, 2 ml of an overnight culture of BL21DE3-pGRO7 in LB containing 12.5 *μ*g/ml chloramphenicol was used to start a 200 ml culture in Zymo Broth (kit # T3002) containing 12.5 *μ*g/ml chloramphenicol. The culture was grown at 300 rpm, 30C until OD_600_ 0.4-0.6 (~4-5 hours) and then immediately plunged into an ice bath with continuous mixing of the culture of the ice for 3 minutes. The flask was left on ice for 7 more minutes, and the centrifuged in 4×50 ml ice-cold sterile tubes at 4 °C for 10 minutes at 2500×g. The supernatant was discarded, and each pellet was resuspended in with 5 ml ice-cold 1X Wash Buffer by gently tapping. The resuspended pellets were then centrifuged at 4 ^°^C for 10 minutes at 2500×g, the supernatant removed, and then each pellet resuspended with 5 ml 1X Competent Buffer. The cells were combined into a single tube and kept on ice until needed.

The 190 PAH protein sequences designed using the ProT-VAE were ordered as gene fragments. The sequences were cloned into a complementary mScarlet-containing expression vector using golden-gate assembly. Briefly, a 10 *μ*L reaction containing 0.5 *μ*L plasmid backbone (100 ng/*μ*L), 3 of the gene fragment, 1 *μ*L of 10× T4 DNA ligase buffer, 0.5 *μ*L of Bsal-HF v2, 0.5 *μ*L T4 DNA Ligase, and 4.5 *μ*L molecular grade water (molar ratio of 2:1 insert:vector) was prepared and then cycled 60 times, keeping the temperature at 37 °C for 5 minutes, and then reducing the temperature to 16 ^°^C for 5 minutes. 2 *μ*L of the assembled plasmids were then added to 40 *μ*l of freshly prepared competent cells which were pre-aliquoted into in an ice-cold Biotix 96-square V-bottom deep well microplate. The plate was sealed with a breathable plate seal and incubated on ice for 10 minutes. Following incubation, 100 *μ*l/well of SOC was added to cells and incubated in an Infors shaker for at 37 °C, 600 RPM, 80% humidity. After a 1 hour incubation 360 *μ*l of LB containing 6.25 *μ*g/ml chloramphenicol and 100 *μ*g/ml carbenicillin was added to each well and incubated in the Infors shaker at 30 °C, 900 RPM, 80% humidity for 20-22 hours.

#### 4.1.3 HT Expression

We make our autoinduction media in-house with the attached protocol. First, 30 *μ*L of the LB-grown transformation mix was added to 570 *μ*L of autoinduction media (25 mM KH2PO4, 47 mM Na2HPO4, 70 mM tryptone, 0.5% yeast extract, 30 mM NaCl, 2.8 mM glucose, 5.84 mM lactose, and 0.6% glycerol) supplemented with 6.25 *μ*g/ml chloramphenicol and 100 *μ*g/ml carbenicillin that had already been aliquoted to each well of a Biotic 96-square V-bottom deep well plate. The plate was covered with an Aera seal and grown for 24 hours at 30 ^°^C, 900 RPM, 80% humidity.

#### 4.1.4 HT Chemical Lysis

Cells were pelleted in expression plates by centrifuging at 4000×g for 30 minutes at 4 °C. The supernatant was decanted then plates were sealed with aluminum plate seals and stored at −80 °C for later use. To make lysis solution Solulyse extraction reagent was supplemented with lysonase (1 *μ*L/ml) and 1 mM Phe, and then 270 *μ*L of this solution was added to each well of the deep well plate to lyse the cells. The plate was covered with a silicone seal and briefly vortexed until all pellets were resuspended. Cells were mixed on an orbital shaker at 900 RPM for 30 minutes at 4 °C and then centrifuged for 30 minutes at 6000×g at 4 °C. 100 *μ*l of lysate was removed and pipetted into a lysate plate for characterization in the HT Lysate Assay.

#### 4.1.5 HT BioAssay

To measure the PAH activity of the lysates we followed the conversion of phenylalanine to tyrosine by measuring the fluorescent emission at 304 nm (ex274nM). Reactions were initiated by transferring 10 *μ*L of the lysate into the appropriate well of a black Greiner *μ*Clear, F-bottom, 96-well microplate and then adding 240 *μ*L of a reaction solution containing 100 mM HEPES pH 7.4, 2 mM DTT, 0.6mg/ml Catalase, 10 *μ*M Fe(II)SO_4_, 75 *μ*M tetrahydrobiopterin (BH4) and variable amounts of Phenylalanine. Progress of the reaction was measured every 40 seconds for 40 minutes and was carried out in technical triplicate. Upon completion of the assay, the amount of PAH variant was determined by measuring the fluorescence of mScarlet (ex569/em593) and comparing the results to a standard curve calculated with in-vitro purified mScarlet-PAH. Velocities were determined by fitting the reaction progress curves to polynomials and taking the max first derivative and kinetic parameters were determined by fitting the reaction velocities to a modified Hill equation. The kcat values were determined by normalizing the Vmax to the amount of enzyme in each well previously determined through mScarlet fluorescence.

## Acknowledgments

We thank Dr. Vedant Sachdeva for assistance in preparing the AlphaFold structural predictions of the synthetic PAH proteins.

## Appendix A Hierarchical organization of SH3 latent space by phylogeny and function

**Fig. A1.**
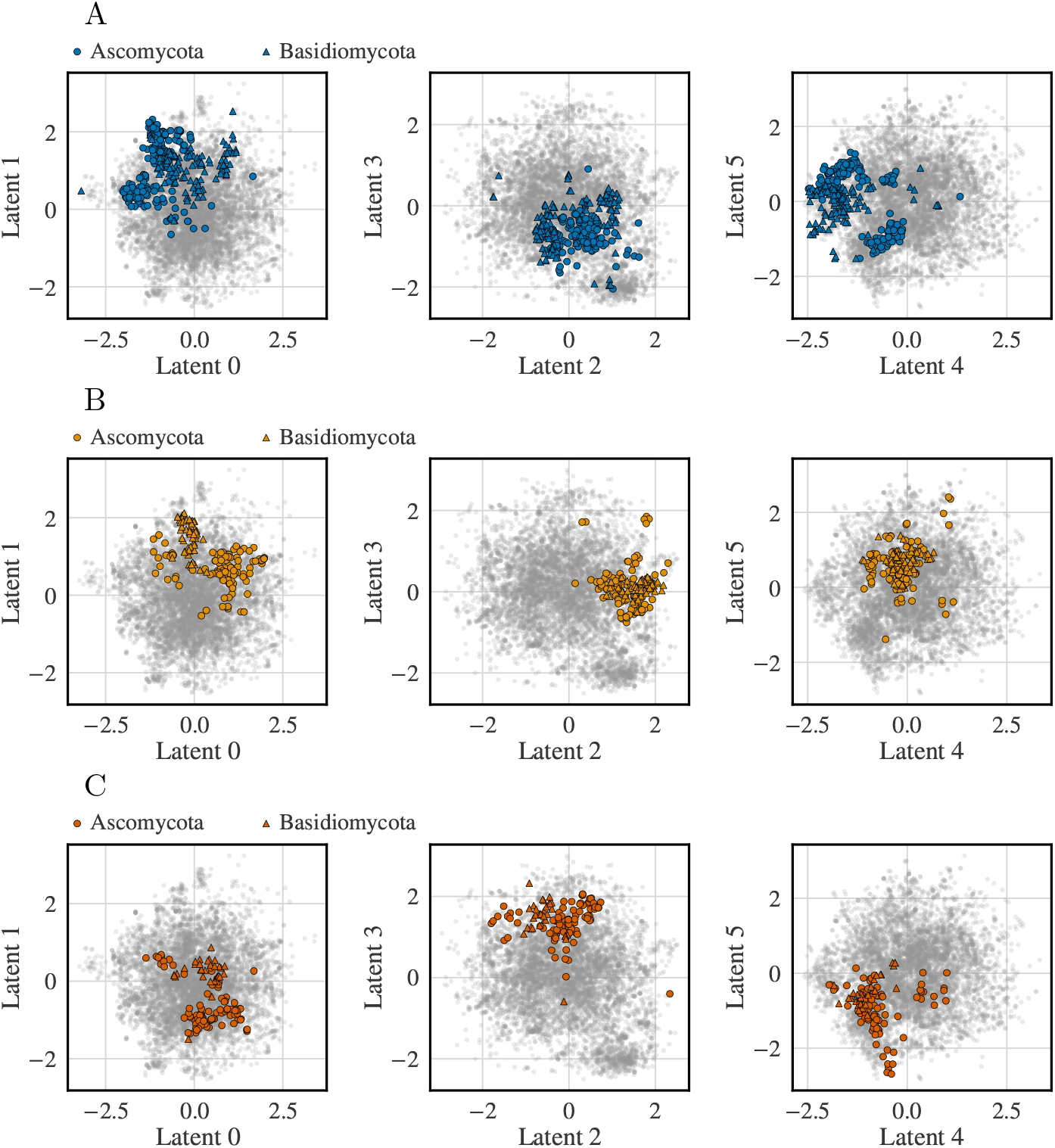
ProT-VAE hierarchically organizes SH3 family first by paralog group and then by phylogeny. A) Two-dimensional projections of the latent space of the Abp1 paralog group separated by phylogeny. B) Two-dimensional projections of the latent space of the Bzz1 paralog group separated by phylogeny. C) Two-dimensional projections of the latent space of the Rvs167 paralog group separated by phylogeny.

